# Dormant Bacteria’s Fatal Attraction to RNA Bacteriophages

**DOI:** 10.64898/2026.04.10.717849

**Authors:** Qiu Zhong, Qianyu Hu, Leilei Wei, Yuhui Yang, Hebin Liao, Zhuojun Zhong, Jiazhen Liu, Xie Fan, Xinyu Jiang, Jianglin Liao, Zongyue Chen, Xuesong He, Liting Wang, Yingying Pu, Jintao Liu, Shuai Le

## Abstract

Phages are the most abundant biological entities on Earth, yet RNA phages are strikingly scarce compared to their DNA counterparts—a long-standing mystery in phage biology. Here, we use dsRNA phage phiYY as a model to demonstrate that while RNA phages efficiently infect growing bacteria, they are gradually eliminated inside dormant bacteria through weak and time-dependent ROS-mediated damage. The RNA phages decline over days rather than through immediate clearance inside dormant bacteria. Accordingly, scavenging ROS with mannitol or overexpressing ROS degradation enzymes AhpB/TrxB2 rescues RNA phages from elimination. Crucially, because the underlying ROS-mediated RNA damage is minimal, RNA phage survival hinges on genomic redundancy. In single-phage infections, the lone RNA genome is highly vulnerable to cumulative damage and is eventually inactivated. In contrast, during co-infection by multiple RNA phages, the presence of multiple genome copies provides functional redundancy, thereby allowing a fraction of RNA phages to survive inside dormant bacteria. Given that most environmental bacteria are dormant and subject to heterogeneous phage infection, this copy number–dependent vulnerability offers a possible explanation for the scarcity, but not perish, of RNA phages in nature.

## Introduction

Bacteriophages (Phages) represent the most abundant biological entities on Earth^1,2^. While tens of thousands of dsDNA phages have been sequenced, RNA phages remain strikingly rare—a long-standing enigma in phage biology^3–7^. In June 2025, we analyzed all completely sequenced viral genomes from the NCBI Viral Genomes Resource. Notably, RNA viruses (excluding over 3 million SARS-CoV-2 sequences) infecting Eukarya and Plants accounted for 57.62% and 53.9% of their respective genomes. In contrast, only 0.53% (131/24,908) of bacterial viruses were RNA phages. None were found to infect Archaea (Fig. 1A). Even the employment of alternative isolation approaches has not been able to significantly increase the identification of RNA phages^8^. Metatranscriptomics studies have revealed a significant increase in both the abundance and diversity of RNA phages. However, their abundance remains limited compared to the vast diversity of DNA phages^9–13^, underscoring the puzzling scarcity of RNA phages in nature^5^.

**Fig. 1:**
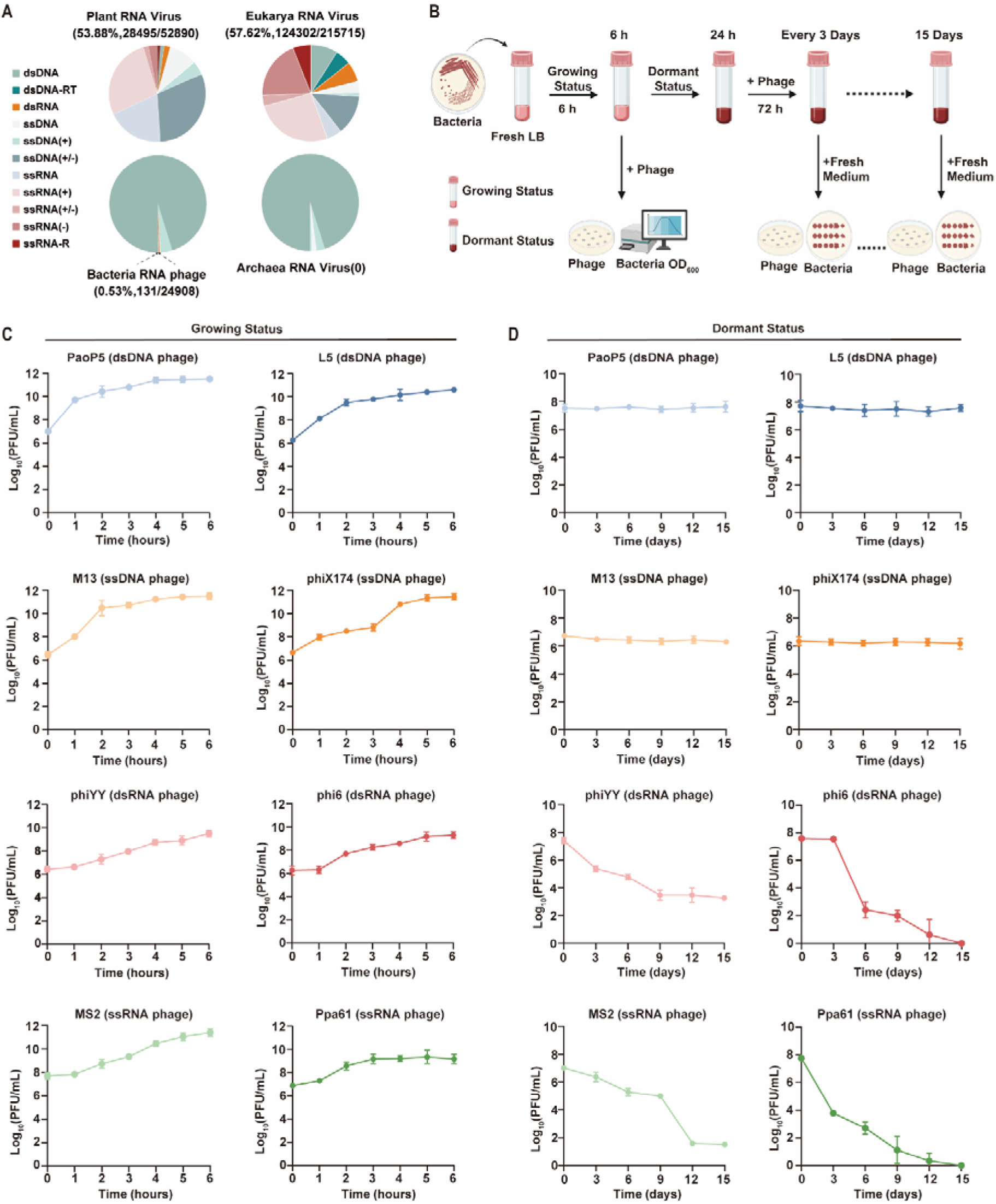
The phage-host dynamics in growing or dormant bacteria. (A) Pie-charts describe the distribution of virus type in Plants, Eukarya, Bacteria, and Archaea. Completely sequenced viral genome data were retrieved from the Viral Genomes Resource (June 2025), excluding over 3 million SARS-CoV-2 sequences. (B) Experimental design schematic: Bacterial cultures were grown in fresh LB medium to early-log phase (OD600=0.2-0.4), then half of the culture was infected with phages (MOI=0.01). Bacterial and phage counts were monitored hourly (0-6 h) to generate growth curves. The remaining half culture was further incubated for 24 h to induce dormancy, followed by phage infection (MOI=0.01) and subsequent 15-day incubation at 37°C. Samples were collected at 3-day intervals for analysis. Bacterial viability was assessed through CFU determination using serial 10-fold dilutions and plating, while phage titers were quantified via efficiency of plating (EOP) assays with fresh host bacteria. All experiments were performed in triplicate. (C) Phage dynamics in log-phase bacteria. Each of the eight phages was used to infect log-phase bacterial cultures, and phage replication was quantified through plaque assays. All phages successfully infected and replicated within their actively growing host bacteria. Three biological repeats were performed for each assay. (D) Phage dynamics in dormant bacteria. Following 24 hours of culturing, bacteria entered a dormant state. Phage titers were monitored at three-day intervals. While all DNA phages remained viable and could be recovered upon addition of fresh medium and host bacteria, RNA phage titers exhibited a continuous decline under the same conditions. Three biological repeats were performed for each assay.

We hypothesize that a universal RNA phage defense mechanism, shared across bacteria, may underlie the rarity of RNA phages. However, known RNA phage defense systems—such as CRISPR-Cas9^14^, leucine-rich repeat-containing proteins (NLRs)^15^, toxin-antitoxin systems^16^, or receptor gene mutations^17^—are highly specific and unlikely to account for the broad suppression of RNA phages.

Most bacteria on Earth exist in a dormant state, exhibiting remarkable resilience to environmental perturbations^18^. While DNA phages can persist by integrating into bacterial genomes^19,20^, the effect of dormancy on RNA phages remains unexplored. To address this, we investigate the survival of eight non-lysogenic phages belonging to four phage types--double-stranded (ds) RNA ((phiYY and phi6), single-stranded(ss) RNA (Ppa61 and MS2), dsDNA (PaoP5 and L5) and ssDNA (M13 and phiX174) phages-- in the presence of their respective growing or dormant bacterial hosts (Table S1).

## Results

Our experimental approach involved subculturing *Pseudomonas aeruginosa*, *Pseudomonas. syringae* or *Escherichia coli* for 24 hours to induce a non-growing, stationary phase^19,21^. To ensure accurate colony-forming unit (CFU) quantification, we used a low phage multiplicity of infection (MOI = 0.01), minimizing phage interference during bacterial quantification (Fig. 1B).

All eight phages successfully replicated in growing host bacteria, with titers increasing continuously throughout the 6-hour experiment (Fig. 1C), though some phages, such as M13 and Ppa61, demonstrated limited lytic activity, allowing concurrent bacterial growth^22^.

Consistent with previous reports ^19,20^, both dsDNA phages (PaoP5 and L5) and ssDNA phages (M13 and phiX174) maintained stable titers for over 15 days in dormant cultures (Fig. 1D). These phages remained viable and could form plaques upon nutrient addition without significant titer loss, demonstrating that phage DNA can persist in dormant bacteria until host regrowth enables replication. The absence of an increasing phage titer indicates proper induction of bacterial dormancy.

In striking contrast, both dsRNA phages (phiYY and phi6) and ssRNA phages (Ppa61 and MS2) exhibited progressive titer declines in dormant cultures, suggesting a significantly weak but continuous RNA phage killing (Fig. 1D). First, to further confirm bacterial dormancy, phiYY and phi6 were incubated with PAO1r or DSM-21482 that had been subcultured for either 24 or 48 hours. During the short-term period (0–8 h), phage titers remained stable, showing no increase or decrease (Fig. S1A–B). To rule out the possibility that the subsequent decline was due to bacterial death, we measured bacterial CFUs. We observed only a modest reduction of at most 10-fold (Fig. S2), far less than the decrease in RNA phage titer, which exceeded a 10,000-fold reduction (Fig. 1D). Taken together, these observations, combined with the previously documented decline over 15 days, confirm that a dormant bacterial state was successfully established. Under these conditions, RNA phages are unable to replicate and are instead eliminated slowly by dormant bacteria.

To determine the site of phage inactivation, we analyzed phage distribution between supernatant (unadsorbed phages) and pelleted bacteria (adsorbed phages), which was resuspended in fresh LB, and phage titer was determined by EOP assay. The rapid decline of RNA phages in the supernatant, coupled with their predominant recovery from bacterial pellets (Fig. 2A), suggests that dormant bacteria mediate RNA phage killing through a post-adsorption mechanism.

**Fig. 2:**
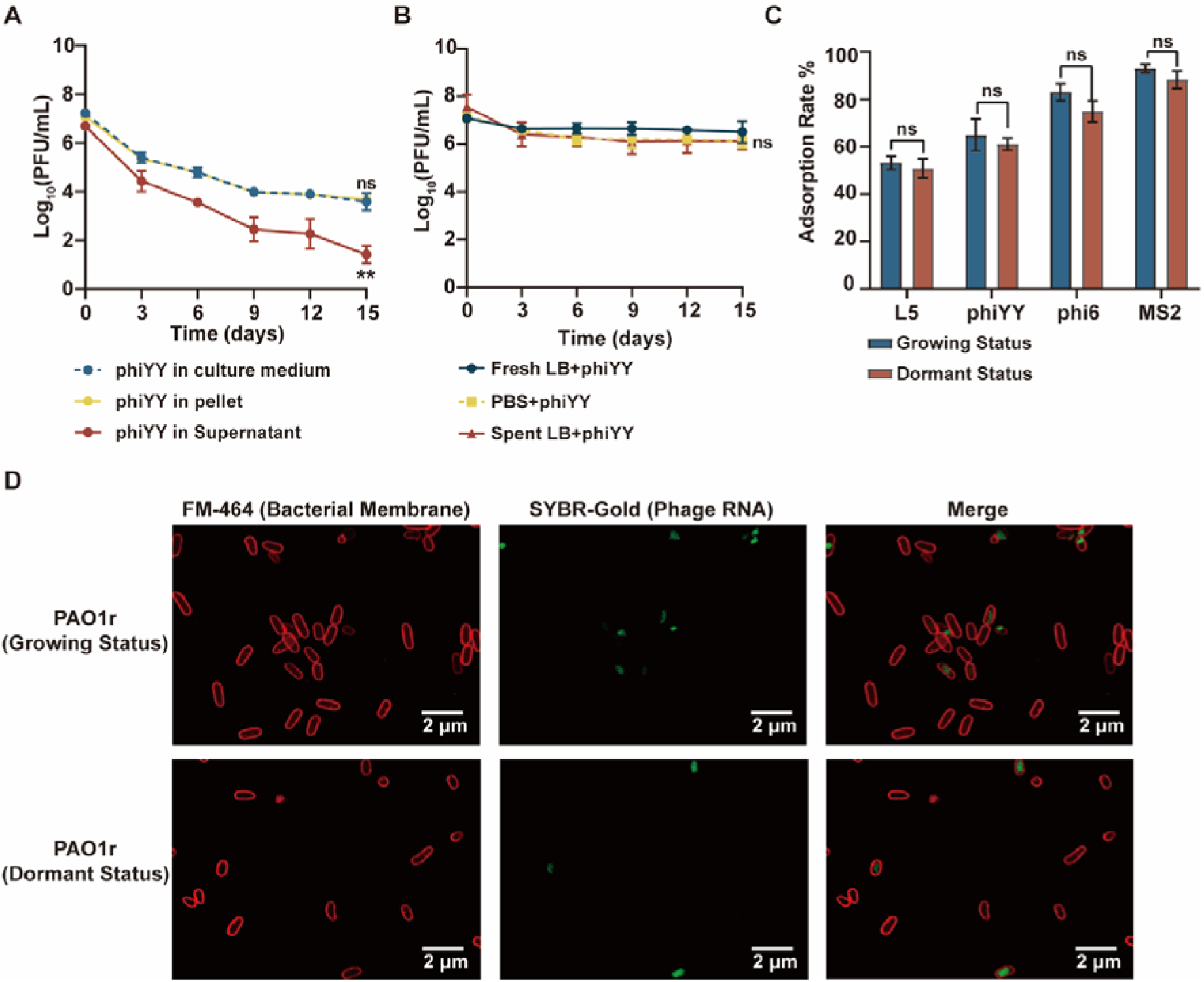
The RNA phage adsorbs and enters the dormant bacteria. (A) Time-course analysis of phage phiYY distribution in whole culture, pellet, and supernatant fractions by EOP assay (n=3). Most phage particles remained cell-associated. Data are presented as mean ± s.d. Two-way ANOVA followed by post hoc test revealed no significant difference between the titer of phiYY in the pellet and that in the culture medium (*P = 0.8744*), whereas the titer in the supernatant was significantly different from that in the culture medium (*P = 0.0019*). *\*\*P < 0.01*; ns, not statistically significant. (B) Phage phiYY was incubated in fresh LB, PBS, or spent medium from 48-hour bacterial cultures in the absence of dormant bacteria. Under all conditions, phage titers remained stable throughout the experimental period. Data are presented as mean ± s.d. (n = 3). Two-way ANOVA followed by post hoc test revealed no significant differences between titers in fresh LB and those in PBS (*P = 0.2544*) or spent medium (*P = 0.1137*). (C) The adsorption rates of phages L5, phiYY, phi6, and MS2 to their respective growing or dormant bacterial hosts. For each phage, adsorption was assessed in three independent biological replicates, and data are presented as mean ± s.d. (n = 3). Statistical significance was determined by the independent samples Student’s t-test comparing adsorption between growing and dormant bacteria. No significant differences were observed for any of the phages tested: L5 (growing: 53.17% ± 2.76% vs. dormant: 50.87% ± 4.05%, *P = 0.6177*); phiYY (growing: 65.03% ± 6.73% vs. dormant: 61.13% ± 2.56%, *P = 0.2720*); phi6 (growing: 82.97% ± 3.59% vs. dormant: 74.87% ± 4.39%, *P = 0.0834*); MS2 (growing: 93.03% ± 1.75% vs. dormant: 88.20% ± 3.65%, *P = 0.2221*). ns, not significant. (D) At 20 minutes post-infection, green fluorescence was observed inside both growing and dormant bacteria, indicating that the RNA phage had successfully entered the host cells in both conditions.

To further verify this, we first assessed phage stability and found that phiYY remains relatively stable in fresh LB, PBS, and spent medium from 48-hour bacterial cultures, the titer of phiYY incubated in fresh LB only decreased form 1.21 ± 0.21×10^7^ PFU/mL to 4.37 ± 3.23 ×10^6^ at day 15 (Fig. 2B), while decreased to 4.80 ± 4.16 ×10^3^ PFU/mL in the presence of PAO1r (Fig. 2A). Second, we examined the adsorption rate of phage L5, phiYY, phi6, and MS2, and observed no significant difference in binding to growing versus dormant bacteria (Fig. 2C). Finally, to determine whether adsorbed phages enter the cells, we performed dual staining with SYBR Gold (phage dsRNA) and FM4-64 (bacterial membrane). We observed their interaction using a Leica Stellaris super-resolution confocal microscope. 20 min after mixing phage with bacteria, green fluorescence was detected inside both growing and dormant bacteria, confirming that phiYY successfully enters dormant cells (Fig 2D). Collectively, these results indicate that the subsequent continuous decrease in phage titer is attributable to intracellular inactivation by the dormant bacteria after entry.

To investigate the molecular mechanism of how dormant bacteria kill RNA phages, we used dsRNA phage phiYY as a model. After adsorbing to the bacteria, phage dsRNA genome and RNA-Dependent RNA Polymerases (RDRP) were transferred into the bacteria^23,24^, which might both be damaged by dormant bacteria. First, we test the impact of RDRP and the dsRNA cleavage enzyme RNaseIII^25^. The overexpression of RDRP or the knockout of RNaseIII does not significantly affect the survival of the dsRNA phage (Fig. 3A), since on day 15, the phiYY titer in PAO1r is not significantly different from that in ΔRNaseIII (P=0.9881) or PAO1r/p-RDRP (P=0.0530). Overexpression of RNaseIII promoted the clearance of phiYY and led to the perish of phiYY at day 15, which indicates that RNaseIII could accelerate the cleavage of dsRNA inside the bacteria and could be one of the factors contributing to RNA phage defense. But the knockout of RNaseIII does not significantly affect it, indicating that without RNaseIII, the RNA genome was still damaged by other factors (Fig. 3A).

**Fig. 3:**
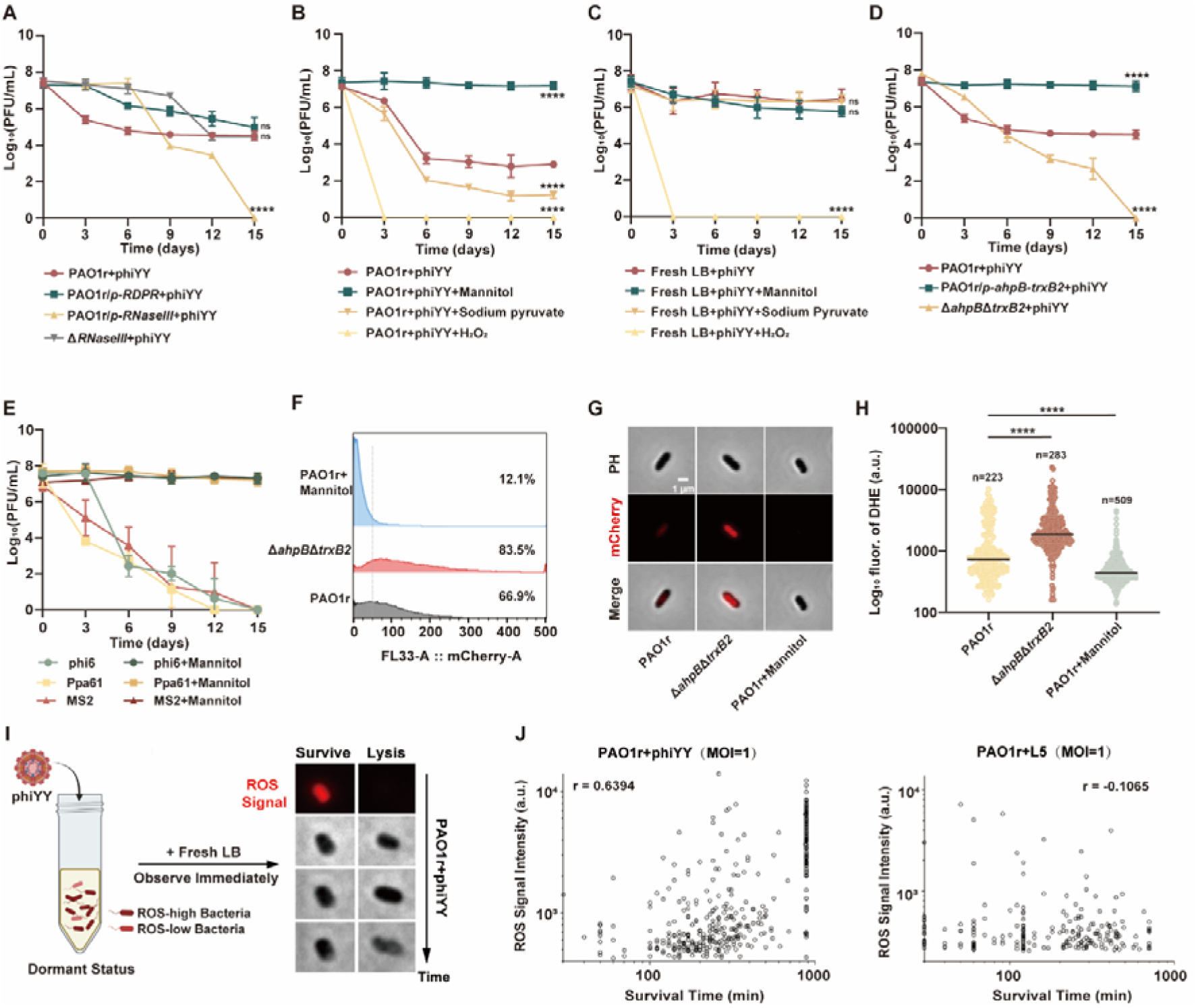
Phage infection dynamics in dormant bacteria with different treatments. (A) Dynamics of phage phiYY in dormant *P. aeruginosa* PAO1r, Δ*RNaseIII*, PAO1r/*p-RDRP*, and PAO1r/*p-RNaseIII*. Data are presented as mean ± s.d. (n = 3 biological replicates). Statistical significance was determined by Two-Way ANOVA with post hoc test, *\*\*\*\*P < 0.0001*, ns, not significant. (B) Dynamics of phage phiYY in dormant PAO1r supplemented with mannitol (230 mM), sodium pyruvate (50 mM), or H_₂_O_₂_ (100 mM). Data are presented as mean ± s.d. (n = 3). Statistical significance between treatment and untreated group was determined by two-way ANOVA followed by a post hoc test. *\*\*\*\*P < 0.0001*. (C) Dynamics of phage phiYY in fresh LB supplemented with mannitol (230 mM), sodium pyruvate (50 mM), or H_₂_O_₂_ (100 mM) in the absence of dormant host bacteria. Data are presented as mean ± s.d. (n = 3). Two-way ANOVA followed by post hoc test revealed no significant differences between the untreated group (phiYY in fresh LB) and phiYY in mannitol-supplemented medium (*P = 0.1365*) or sodium pyruvate-supplemented medium (*P = 0.9286*). In contrast, phiYY in H_₂_O_₂_-supplemented medium showed a highly significant reduction compared to the untreated group (*P < 0.0001*). Statistical significance relative to the untreated control is indicated as follows: *\*P < 0.05*, *\*\*P < 0.01*, *\*\*\*P < 0.001*, *\*\*\*\*P < 0.0001*; ns, not significant. (D) Dynamics of phage phiYY in dormant *P. aeruginosa* PAO1r, Δ*ahpB-trxB2*, PAO1r/*p-ahpB-trxB2*, and PAO1r/*p-RNaseIII*. Data are presented as mean ± s.d. from three independent biological replicates. Statistical significance was determined by two-way ANOVA followed by post hoc test to compare with the control group. *\*\*\*\*P < 0.0001*. (E) Mannitol supplementation significantly stabilized three other RNA phages—including the dsRNA phage phi6 and the ssRNA phages Ppa61 and MS2, in co-culture with their dormant host bacteria over 15 days. (F) Flow cytometry analysis of intracellular ROS in 72 h-cultured dormant bacteria (mCherry^+^ threshold = 50 a.u., dashed line). (G) Representative DHE-stained images of D3-phase bacteria: PAO1r (left), Δ*ahpB*Δ*trxB2* (middle), and PAO1r+mannitol (right). Scale bar = 1 μm. (H) Single-cell ROS quantification of intracellular ROS levels. (median shown in black; PAO1r: n=223; Δ*ahpB*Δ*trxB2*: n=283; PAO1r+mannitol: n=509. Statistical significance was determined using One-Way ANOVA with Dunnett’s multiple comparisons test. For PAO1r and Δ*ahpB*Δ*trxB2*, *P* = 5.469*10^-10^; for PAO1r and mannitol, *P* = 3.048*10^-13^; for Δ*ahpB*Δ*trxB2* and mannitol treatment, *P* = 9.152*10^-32^). (I) Representative images illustrating the fate of dormant bacteria following phage infection. We hypothesize that bacteria with high intracellular ROS levels (left) survive phage infection, whereas those with low intracellular ROS (right) undergo lysis. (J) Correlation between bacterial survival time and intracellular ROS level in dormant bacteria infected with dsRNA phage phiYY or dsDNA phage L5 (MOI=1). No correlation was observed between the ROS level and the bacteria survival time as analyzed using the Pearson correlation coefficient (left, *r* = 0.1226, right, *r* = -0.1065).

Then we hypothesized that the intracellular reactive oxygen species (ROS) might damage the RNA because the ROS is common in all bacteria, which could damage RNA, DNA, and protein^26^. Bacteria have multiple DNA repair systems but lack RNA repair systems; thus, the ROS-induced RNA damage might be lethal to RNA phages^26^, while DNA phages could survive the ROS-induced DNA damage due to the bacterial repair systems.

To test this hypothesis, we first added mannitol (OH- scavenger)^27^ or sodium pyruvate (H_2_O_2_ scavenger)^28^ into the medium, and, interestingly, phiYY could co-exist with a dormant host and survive for 15 days without significant decline in the presence of mannitol but not sodium pyruvate, and the clearance of RNA phage was accelerated by the supplement of H_₂_O_₂_ (Fig. 3B). To rule out any direct effect of these compounds on the phages, we incubated phiYY in culture medium supplemented with mannitol, sodium pyruvate, or H_₂_O_₂_ in the absence of bacterial hosts. Final concentrations of 230 mM mannitol and 50 mM sodium pyruvate had no impact on phage stability, whereas 100 mM H_₂_O_₂_ rapidly inactivated the phages (Fig. 3C). To further rule out the possibility that mannitol supplementation might promote phage replication by reactivating bacterial growth, we tested its effect on dormant bacteria and found that supplementation of mannitol to 230 mM did not enhance phage replication over 8 hours, indicating that bacterial dormancy was not disrupted (Fig. S1C). These control experiments further support that the protective effect of mannitol on phiYY occurs intracellularly, rather than through direct stabilization of the phage particle (Fig. 3C).

The genetic data are consistent with this; knocking out the alkyl hydroperoxide reductase (*ahpB*) and thioredoxin reductase 2 (*trxB2*), which degrade ·OH^29,30^, significantly promoted the death of RNA phages, and all RNA phages are cleared by day 15. In contrast, overexpression of *ahpB-trxB2* promoted the survival of RNA phage, with titers remaining at 1.53 ± 1.10 ×10^7^ PFU/mL at day 15, showing no significant decrease compared to the initial titer of 2.27 ± 0.83 ×10^7^ PFU/mL (Fig. 3D). Moreover, the addition of mannitol also promoted the survival of the other RNA phages included in the study; phi6, Ppa61 and MS2 (Fig. 3E). These data indicate that intracellular ROS could kill RNA phages inside the dormant bacteria, which leads to the death of RNA phages, while the cleavage of ROS inside bacteria, using mannitol or overexpression of *ahpB-trxB2,* could maintain the survival of RNA phages.

To elucidate the role of ROS in RNA phage clearance, we first assessed intracellular ROS levels at the population level using dihydroethidium (DHE), a cell-permeable fluorescent probe that emits red fluorescence upon ROS oxidation. After 72 hours of cultivation, flow cytometry revealed that 83.5% of Δ*ahpB*Δ*trxB2* mutant cells were mCherry-positive, showing significantly elevated ROS levels compared to wild-type PAO1r (66.9%). Notably, PAO1r cells treated with the ROS scavenger mannitol exhibited the most pronounced ROS reduction, with only 12.1% displaying mCherry fluorescence (Fig. 3F). These population-level findings were corroborated by single-cell fluorescence microscopy, which consistently showed higher ROS accumulation in the mutant strain versus both wild-type and mannitol-treated cells (Fig. 3G-H).

Because ROS impacts RNA phage survival, and the bacterial culture has a heterogeneity in ROS levels, we initially hypothesized that this ROS heterogeneity contributes to phages surviving in bacteria with low ROS but are eliminated in those with high ROS (Fig. 3I).

To directly examine ROS-phage interactions, we performed real-time fluorescence microscopy on DHE-stained bacterial cultures (72-hour growth) during phage infection. We hypothesized that ROS-high bacteria would resist RNA phage infection while ROS-low bacteria would be susceptible. Indeed, several ROS-high bacteria remained intact for ≥12 hours post-infection, whereas ROS-negative bacteria lysed within 5 hours (Fig. 3I). However, quantitative analysis showed no significant correlation between intracellular ROS intensity and survival time during phiYY infection (MOI=1) (Fig. 3J). And ROS levels showed no correlation with survival following DNA phage L5 infection (Fig. 3J). This aligns with our growth curve data showing gradual RNA phage clearance over days (Fig. 1D), not immediately (Fig. S1A), likely because excessively high ROS would be cytotoxic and bacterial ROS-scavenging enzymes tightly regulate concentrations^26^, thus even the high-ROS we observed is not high enough to directly kill RNA phage, but could kill it through a time-dependent accumulation of minimal RNA damage.

To test our hypothesis, we first sought to use real-time monitoring. However, given the inability to sustain fluorescent staining over multiple days, we designed an alternative controlled infection assay. We compared infection outcomes in wild-type PAO1r, the high-ROS mutant Δ*ahpB*Δ*trxB2*, and mannitol-treated PAO1r with suppressed ROS (ROS-low) across different growth phases (Fig. 4A).

**Fig. 4:**
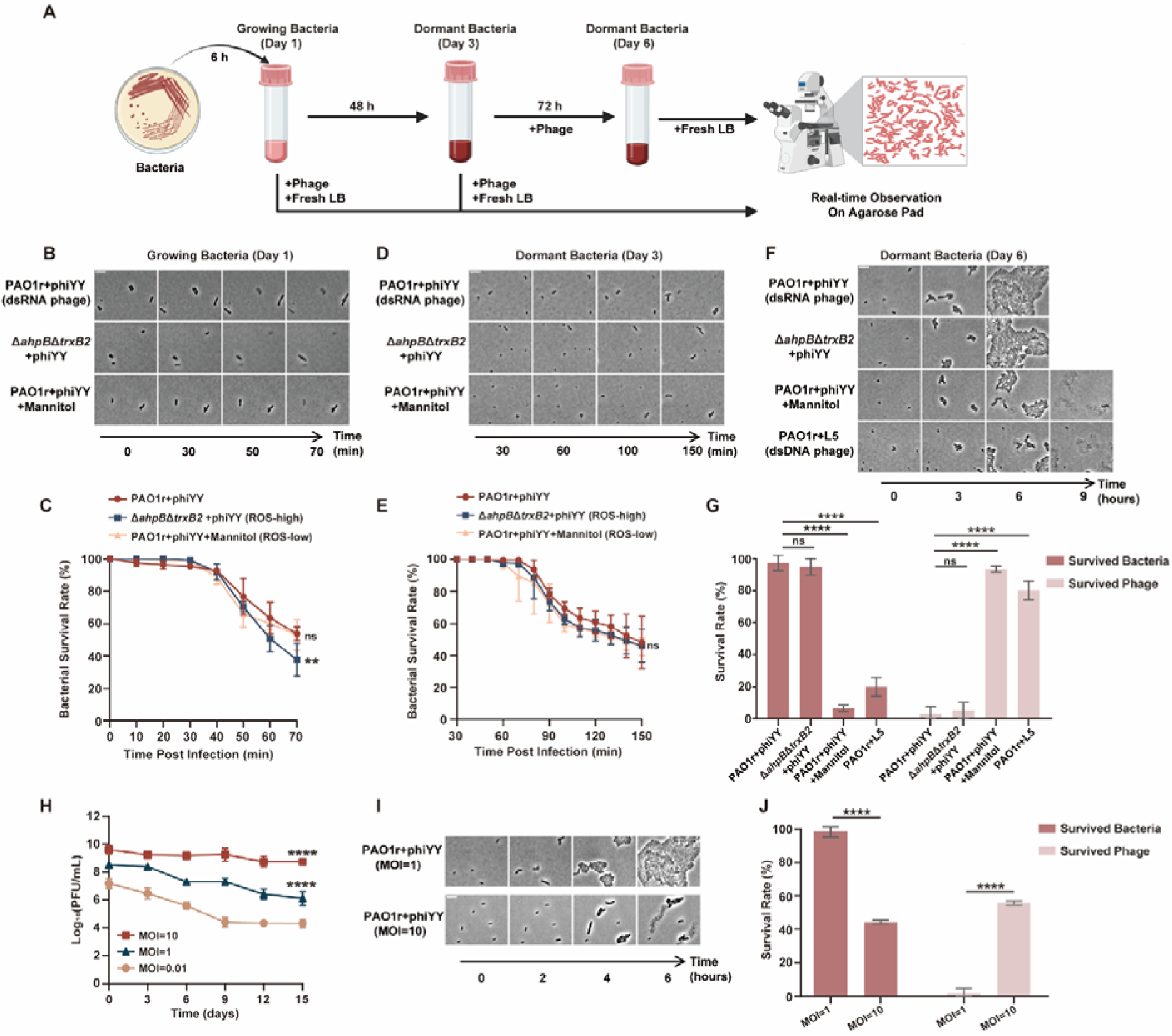
Outcome of phages infecting growing or dormant bacteria visualized in the microscope. (A) Experimental design schematic: Bacterial cultures were grown in fresh LB medium to early-log phase (OD600=0.2-0.4), then part of the log phase bacteria (Day 1) was infected with phages, and the growth of bacteria was visualized in microscopy. Then the bacteria were further cultured for a total of 48h to induce a deep dormancy (Day 3). At this point, the dormant bacteria were mixed with phage. One part of the mixture was transferred to a fresh LB agarose pad for microscopic observation; the other part was cultured for an additional 3 days (Day 6) after which the cells were transferred to a fresh LB agarose pad, and bacterial survival was assessed by microscopy. (B) Representative phase-contrast images showing the progression of phage phiYY infection (MOI = 1) in growing cells (Day 1) on LB agarose pads. From top to bottom: wild-type PAO1r, Δ*ahpB*Δ*trxB2* mutant, and PAO1r supplemented with mannitol. Scale bar, 1 μm. (C) Single-cell fates were continuously monitored at 10-min intervals for 70 min post-infection. Bacterial survival rates were quantified and are presented as mean ± s.d. from three independent experiments. Statistical significance was determined by two-way ANOVA followed by post hoc test. The survival rate of PAO1r treated with mannitol prior to phiYY infection was significantly different from that of the untreated control group (*P = 0.0092*). *\*\*P < 0.01*; ns, not significant. (D) Representative phase-contrast images showing the progression of phage phiYY infection (MOI = 1) in dormant cells (Day 3) on LB agarose pads. Cells were incubated with phiYY in the original medium for 30 min, then spotted onto fresh LB agarose pads for time-lapse imaging. Single cells were monitored for 150 min at 10-min intervals to capture infection outcomes during resuscitation. Scale bar, 1 μm. And the statistics of bacterial survival rate are shown in (E). (F) Representative phase-contrast images showing bacterial survival status after 3-day co-culture with phage phiYY (MOI = 1) on LB agarose pads. Dormant cells (D3) were infected with phiYY and co-cultured for another 3 days prior to imaging. The mixture was spotted onto fresh LB agarose pads to induce resuscitation. Single-cell fates were monitored at 15-min intervals for 9 hours. Scale bar, 1 μm. (G) Quantification of surviving and lysed bacteria (the latter indicating successful phage infection and replication) at 9 hours post-transfer to fresh culture medium. Data are presented as mean ± s.d. (n = 3). For each group, bacterial fates were monitored by time-lapse microscopy, with 50 randomly selected initial cells tracked per replicate. (H) Dynamics of phage phiYY in dormant *P. aeruginosa* PAO1r with an MOI of 0.01, 1 and 10, respectively. Data are presented as mean ± s.d. (n = 3). Statistical significance was determined by a two-way ANOVA post hoc test. Statistical significance compared to the control group (MOI=0.01) is indicated: *\*\*\*\*P < 0.0001*. (I) Representative phase-contrast images showing bacterial survival status after 3-day co-culture with phage phiYY at MOI 1 or 10 on LB agarose pads. Dormant cells (D3) were infected with phiYY and co-cultured for 3 days prior to imaging. The mixture was spotted onto fresh LB agarose pads to induce resuscitation. Single-cell fates were monitored at 15-min intervals for 6 hours. Scale bar, 1 μm. (J) Dormant cells were (D3) infected with phage phiYY at MOI 1 or 10 and co-cultured for 3 days, then the surviving and lysed bacteria (the latter indicating successful phage infection and replication) were quantified 9 hours post-transfer to fresh culture medium. Data are presented as mean ± s.d. (n = 3). Statistical significance was determined by a two-way ANOVA post hoc test. Statistical significance compared to the control group (MOI=1) is indicated: *\*\*\*\*P < 0.0001*.

During logarithmic (D1) and dormant (D3) phases, phiYY infection (MOI=1) resulted in similar mortality rates (∼50%) within 70 min (D1) or 150 min (D3) (Fig. 4B-E) upon transfer to a fresh LB agarose pad. To rule out secondary infection effects, enumeration time points were selected based on the division time of uninfected controls (70 min for D1, 150 min for D3). The comparable mortality at these early time points suggests that weak, ROS-mediated killing is not effective in eliminating RNA phage in growing or nutrient-amended dormant bacteria.

After three days of co-incubation (D6), striking differences emerged. In wild-type cultures, only 2.78% ± 4.81% lysed bacterium was observed (Fig. 4F-G). In contrast, mannitol-treated (ROS-low) bacteria showed 93.39% ± 1.83% lysis. This demonstrates that intracellular ROS inactivates phiYY, while ROS scavenging promotes phage survival. As a control, the dsDNA phage L5 lysed 80.05% ± 5.87% of wild-type bacteria after three days of co-incubation (D6), confirming that ROS doesn’t affect DNA phages. These data confirm that ROS-mediated damage to the RNA phage genome is extremely weak and time-dependent.

The final question is how such ROS damage eliminates phage RNA without severely affecting bacterial mRNA. We propose that bacterial antioxidant enzymes tightly control ROS at a low, safe level, resulting in minimal RNA damage that accumulates over days, and bacteria have multiple mechanisms to maintain mRNA stability, and damaged bacterial mRNA is replaced through low-level transcriptional activity during dormancy. In our standard low-MOI (0.01) model, most bacteria are infected with a single phage particle containing only one genomic RNA copy, making it highly vulnerable to this cumulative damage. In contrast, the abundant, multi-copy pool of bacterial mRNAs can tolerate sporadic losses without lethal consequences.

To test this hypothesis, we infected bacteria at a high MOI of 10. In striking contrast to the low-MOI scenario, phage survival increased dramatically, with titers remaining at 5.83 ± 2.52 ×10^8^ PFU/mL at day 15, showing a weak decrease compared to the initial titer of 4.67±3.40 ×10^9^ PFU/mL (Fig. 4H). This was further visualized at the single-cell level by microscopy. By day 6, single-cell analysis revealed that the phage survival rate increased from 1.72% ± 2.44% in the low-MOI group (MOI = 1) to 55.71% ± 1.24% in the high-MOI group (MOI = 10) (Fig. 4I–J), consistent with population-level survival curves (Fig. 4H). In summary, our findings indicate that ROS-dependent killing of RNA phages is a weak, time-dependent process.

## Discussion

Our study reviewed a very intriguing impact of bacterial growth status on RNA phage: while RNA phages efficiently kill growing bacteria, dormant bacteria progressively inactivate RNA phages through a slow, ROS-dependent mechanism (FIG 5). Using dsRNA phage phiYY as a model, we showed that intracellular ROS induces cumulative damage to RNA phage genomes in a time-dependent way, requiring several days to achieve inactivation, a process only feasible in dormant bacteria. This subtle phenotypic difference was validated through both genetic analysis and single-cell visualization, which illuminates bacterial dormancy as an underappreciated defense strategy, transforming dormant bacteria into effective predators of RNA phages. Current studies of phage defense mechanisms are mostly investigated in growing cultures^31–33^, further investigation into crosstalk between dormancy and canonical phage defense systems (e.g., CRISPR, restriction-modification) may reveal more interesting different principles of phage-bacteria interactions.

**Fig 5:**
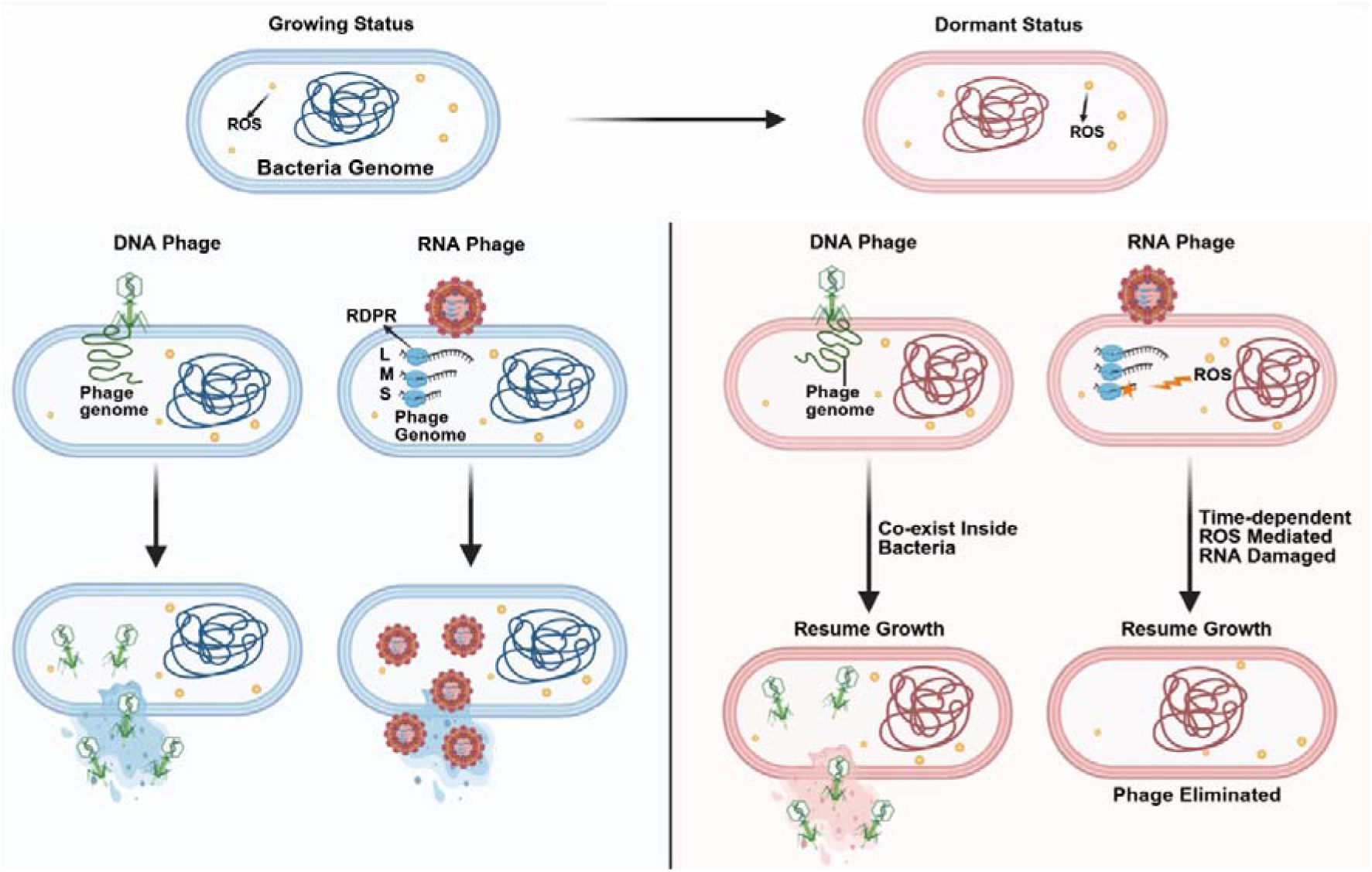
Dormant bacteria kill RNA phage. Both RNA and DNA phages kill growing bacteria. However, when bacteria become dormant, RNA phages co-localize within bacteria but undergo time-dependent accumulation of ROS-mediated RNA damage (orange star lightning), leading to a slow loss of recoverable phage. On the contrary, DNA phages remain stable intracellularly, maintaining infectivity until host resuscitation.

The scarcity of RNA phages is a longstanding mystery in phage biology^5^. Previous efforts to detect RNA phages using diverse approaches have suggested that this scarcity reflects a genuine ecological phenomenon rather than a consequence of limited isolation^5,12^. Given the prevalence of bacterial dormancy in natural environments^18,34^, we propose that the elimination of RNA phages by dormant bacteria may contribute substantially to their low abundance in the wild. This ecological hypothesis now warrants direct validation through field studies and in situ environmental sampling, which will be critical to fully elucidate the mechanisms underlying RNA phage scarcity.

Nevertheless, RNA phages, though rare, have not gone extinct in nature^35^. This persistence can be explained by the modest nature of ROS-mediated RNA damage: it is sufficient to gradually inactivate infecting phages, yet mild enough that a fraction of RNA phages can still transiently survive within dormant bacteria (Fig. 1D). More importantly, this damage is sufficiently weak that co-infection of a dormant bacterium with multiple RNA phage genomes substantially increases phage survival (Fig. 4I-J). Dormant bacteria infected with a single phage particle are likely to eliminate it through cumulative ROS-mediated RNA damage, whereas those infected with multiple copies can preserve at least a subset of the phage population. Such genomic redundancy thus provides a plausible mechanism preventing the complete extinction of RNA phages in natural environments, allowing a subset of phages to persist through prolonged host dormancy. In contrast, DNA phages maintain infectivity during bacterial dormancy through dedicated latency mechanisms and reactivate upon host resuscitation, a strategy that confers a significant ecological advantage^20^.

This finding raises a compelling evolutionary question: Why do RNA viruses thrive in mammals, whereas RNA phages remain scarce in bacterial hosts (Fig 1A)? We propose that this disparity might stem from fundamental differences in host physiology. Unlike bacteria, most mammalian cells do not enter deep dormant states^36^, which limits prolonged intracellular ROS exposure to RNA viruses, thus might facilitate the survival of RNA viruses.

Clinically, phage therapy holds promise as a promising approach to combat antibiotic-resistant bacteria^37^, while this study might explain the limited efficacy of RNA phage therapy^38,39^. In prior clinical trials with chronic *P. aeruginosa* infections, RNA phage phiYY suppressed but failed to eradicate bacterial burdens. We posit that those dormant bacterial subpopulations, which could gradually inactivate phages via ROS, underpinned this therapeutic failure. Thus, targeting dormant reservoirs represents a critical frontier for improving phage therapy outcomes.

While our study demonstrates that dormant bacteria can eliminate RNA phages via ROS, several key questions remain unanswered. First, due to the limitations of dye-based staining methods, we were unable to perform single-cell and real-time analyses of the dynamic interactions among phage infection, ROS heterogeneity, and bacterial viability. Second, the specific ROS species responsible for phage RNA damage, as well as the precise molecular mechanisms leading to phage inactivation, remain unidentified. Finally, field studies investigating the impact of bacterial dormancy on RNA phage ecology represent a critical next step. Future research should address these gaps by employing advanced single-cell tracking techniques, detailed molecular characterization of ROS-induced RNA damage, and in situ environmental sampling to fully elucidate this phenomenon.

## Methods

### Bacterial strains, phages, and culture conditions

The bacterial strains and phages in this work are listed in Table S1. *P. aeruginosa* or *E.coli* strains were grown in Lysogeny Broth (LB) medium at 37°C with 220 r.p.m. shaking. *Pseudomonas syringae* were grown in Lysogeny Broth (LB) medium at 28°C with 220 r.p.m. shaking.

### Induction of Bacterial Dormancy and Phage Co-culture

To induce dormancy, a single bacterial colony was inoculated into 3 mL of LB broth and cultured at 37°C with shaking at 220 rpm for 24 hours, achieving a dormant state. Subsequently, the corresponding phage was added to the bacterial culture for co-incubation at 37°C with shaking at 220 rpm for a given time. Mannitol, H_2_O_2,_ or sodium pyruvate was added to a final concentration of 0.3 M, 10mM, and 50 mM, respectively. On day 7, an additional 1.5 mL of 1 M mannitol was supplemented to the 5 mL culture to maintain its concentration. *P. syringae* and phage phi6 were cultured and tittered at 28 °C.

Phage titration by double-layer agar assay. Every 3 days during co-culture, phage titers were quantified using the double-layer agar method^40^. Briefly, the culture was serially diluted 10-fold in sterile medium, aliquots (10 μL) of each dilution were mixed with log-phase host bacteria, the mixture was mixed with soft agar (0.4% agar) and overlaid onto solid LB agar plates, which were incubated statically at 37°C overnight, then the plaque-forming units (PFU) were counted to calculate phage titers. Bacterial counts were determined by performing 10-fold serial dilutions of the culture, and plating aliquots (100 μL) onto solid LB agar plates, which were incubated statically at 37°C overnight, then counting colony-forming units (CFU). The co-evolution dynamics between phage and bacteria were analyzed by plotting phage titers (PFU/mL) against bacterial counts (CFU/mL) over time.

Phage stability was assessed by incubating phages in fresh LB, PBS, spent medium from 48-hour bacterial cultures for 15 days, mannitol (0.23 M), and sodium pyruvate (50 mM), H_2_O_2_ (50 mM), with phage titers monitored at indicated time points. Spent medium was prepared by pelleting and filtering 48-hour bacterial cultures, followed by the collection of the supernatant.

### Construction of the gene knockout or overexpression bacterial strains

The mutant strains were constructed using homologous recombination to knockout *ahpB/trxB2* and RNaseIII genes in PAO1r as described previously^41^. For *ahpB/trxB2* knockout, the upstream and downstream fragments were amplified by PCR using primer pairs LA-F/R and RA-F/R (Supplementary Table 2), the gentamicin resistant gene was amplified by PCR using primer pairs Gm-F/R, the three fragments were ligated into pEX18Tc vector using the Gibson assembly method to generate pEX18Tc-*ahpB/trxB2*, which was then electroporated into PAO1r competent cells and plated on LB agar containing 20 μg/mL gentamicin, followed by selection of gentamicin-sensitive and sucrose-resistant colonies on LB agar with 320 μg/mL gentamicin and 15% sucrose, with successful *ahpB/trxB2* deletion confirmed by PCR screening.

Knockout of RNaseIII. The upstream fragment (519 bp) and the downstream fragment (481 bp) of the RNaseIII gene was amplified by PCR using the primer pair LA-F/R and RA-F/R. The two PCR fragments were ligated into the expression vector pPAGML via Gibson assembly and transformed into *E. coli* S17-1λ competent cells. The plasmid pPAGML-RNaseIII was transferred into PAO1r through conjugation and plated on a medium containing 20 μg/mL gentamicin (Gm), 50 μg/μL kanamycin (Kan), 100 μg/mL ampicillin (Amp) for overnight selection. After culturing at 37°C until single colonies grew, the single colonies were picked and inoculated into LB liquid medium containing 300 μg/mL Amp until turbidity was observed. Then, they were transferred to an LB plate supplemented with 0.1% PCPA (4-Chloro-DL-phenylalanine) for three-line streaking. Single colonies were picked for PCR verification to check for the absence of RNaseIII, and finally, the Δ*RNaseIII* mutant strain was obtained.

For overexpression strain construction, the *ahpB-trxB2* gene fragment was amplified using ahpB-trxB2-F/R primers, cloned into linearized pHERD20T vector to generate p-*ahpB-trxB2*, electroporated into PAO1r, selected on gentamicin-containing plates, and induced with 0.2% arabinose for gene expression. Other strains were constructed with the same methods using the primers listed in TABLE S2.

### Flow cytometry for intracellular ROS quantification

For the analysis of intracellular ROS levels in the bulk system, we incubated bacteria at various stages with 50 μM dihydroethidium (Aladdin) at 37 °C for 30 min. Then we diluted these samples 100 times in PBS and immediately analyzed them with BD FACS Calibur flow cytometry (Becton Dickinson) using PE-Texas Red channel, a 561-nm green-to-yellow laser, plus a 600 nm long pass filter and a 610/20 nm band-pass filter. For each sample, 10000 events were measured.

### Measurement of intracellular ROS at single cell level

To visualize the intracellular ROS at single cell level, we incubated bacteria with 50 μM dihydroethidium at 37 °C for 30 min. Then we dropped 10 μl of the mixture on a glass slide and covered it with a coverslip, and samples were observed with phase contrast and fluorescent microscopy. The microscopy used was Olympus IX83 (Japan) with Andor’s Zyla 4.2 sCMOS camera (UK) and 100X objective lens.

### Phage interaction with single cells

To explore the relationship between the fate of cells after phage infection and intracellular ROS levels, we incubated the 72-hour-old bacterial culture with 50 μM dihydroethidium at 37 °C for 30 min, then added phage at different MOI.

After diluting the mixture 100 times in PBS, we dropped 10 μl of the mixture over fresh LB agarose pads immediately and CellSens Dimension software (Olympus, version 1.18) was used to monitor the dynamics of bacteria, and images were taken every 10 minutes. The microscope settings were the consistent with those described above.

### Confocal Microscopy

To visualize phage entry into growing and dormant *P. aeruginosa* PAO1r. Dormant bacteria were obtained by incubating a single colony at 37°C for 48 hours. Then, both growing and dormant cultures were adjusted to an optical density (OD_₆₀₀_) of 0.3–0.4. Phage phiYY was stained with SYBR Gold (Thermo Fisher) at a 1:10,000 dilution for 20 minutes^42^. Then, 1 mL of each bacterial suspension was mixed with the stained phages at a multiplicity of infection (MOI) of 10. At 20 minutes post-infection, samples were collected and centrifuged at 5,000 × g for 1 minute. The bacterial pellets were resuspended in 50 µL of phosphate-buffered saline (PBS). For membrane staining, the resuspended cells were incubated with 57 µM FM4-64 membrane dye (Thermo Fisher). Then, one microliter of the stained suspension was placed on a slide and observed under a Leica Stellaris super-resolution microscope (Leica Microsystems). Fluorescence signals were detected using photomultiplier tube detectors. The LIGHTNING real-time super-resolution processing module was activated to enhance image resolution.

### Assessment of bacterial survival following phage infection by microscopy

To assess bacterial survival following phage infection at the single-cell level, sample preparations were performed under three distinct conditions before microscopic examination: 1) Log-phase bacteria: cultures were infected with phage phiYY at MOI 1. Following a 5-min adsorption period at room temperature, the mixture was spotted onto an LB agarose pad. 2) Dormant (48-hour) cultures: phage phiYY (MOI 1) was added to 48-hour stationary-phase cultures. The mixture was incubated at 37 °C with shaking (220 rpm) for 30 min in the original medium, and subsequently transferred onto a fresh LB agarose pad to induce resuscitation. 3) Prolonged co-cultures: bacterial cultures grown for 48 hours were harvested and diluted 100-fold in their own sterile-filtered supernatant to maintain starvation conditions. Phage phiYY was then introduced (MOI 1 or 10), and the co-culture was incubated at 37°C with shaking (220 rpm) for 3 days. Following this prolonged incubation, samples were directly transferred onto a fresh LB agarose pad for microscopic examination.

### Image analysis

Data analysis was performed using ImageJ (National Institutes of Health, USA) and MATLAB (MathWorks, USA). First, individual frames were initially saved in the proprietary .vsi format (default file format of the CellSens Dimension software) and these .vsi files were converted to the standard .tiff format using ImageJ (1.53q). The phase contrast images of cells were segmented using a custom Omnipose model based on MATLAB^43^. The single-cell growth and lysis traces were tracked over time using features extracted at each frame.

## Statistical analysis

Statistical analysis was performed using Student’s *t*-test for comparisons between two groups. One-Way ANOVA was used to compare differences among three or more independent groups, while Two-Way ANOVA was applied to analyze the effects of two independent variables and their potential interaction on the dependent variable. A significance level of *P< 0.05* was adopted to determine statistical significance.

## Acknowledgments

This study was supported by grants from the National Key Research and Development Program of China (2021YFA0911200 to S.L; 2023YFC2306300 to J.L), the National Natural Science Foundation of China (32570191 to S.L, 32170099 to J.L), New Chongqing Youth Innovation Talent Project (CSTB2025YITP-QCRCX0026 to Q.Z), and Chongqing Young and Middle-aged High-level Medical Talents (YXGD202566 to Q.Z).

## Author contributions

Conceptualization, Shuai Le;

Methodology, Zhong Qiu, Hu Qianyu, Leilei Wei, Yuhui Yang, Liao Hebing, Zhong Zhuojun, Pu Yingying, Xie Fan, Liting Wang;

Validation, Xingyu Jiang, Jianglin liao;

Investigation, Zhong Qiu, Hu Qianyu, Leilei Wei, Yuhui Yang;

Data curation, Leilei Wei, Jiazhen Liu ;

Writing – review & editing, Shuai Le, Xuesong He, Jintao Liu;

---

**TABLE S1:**
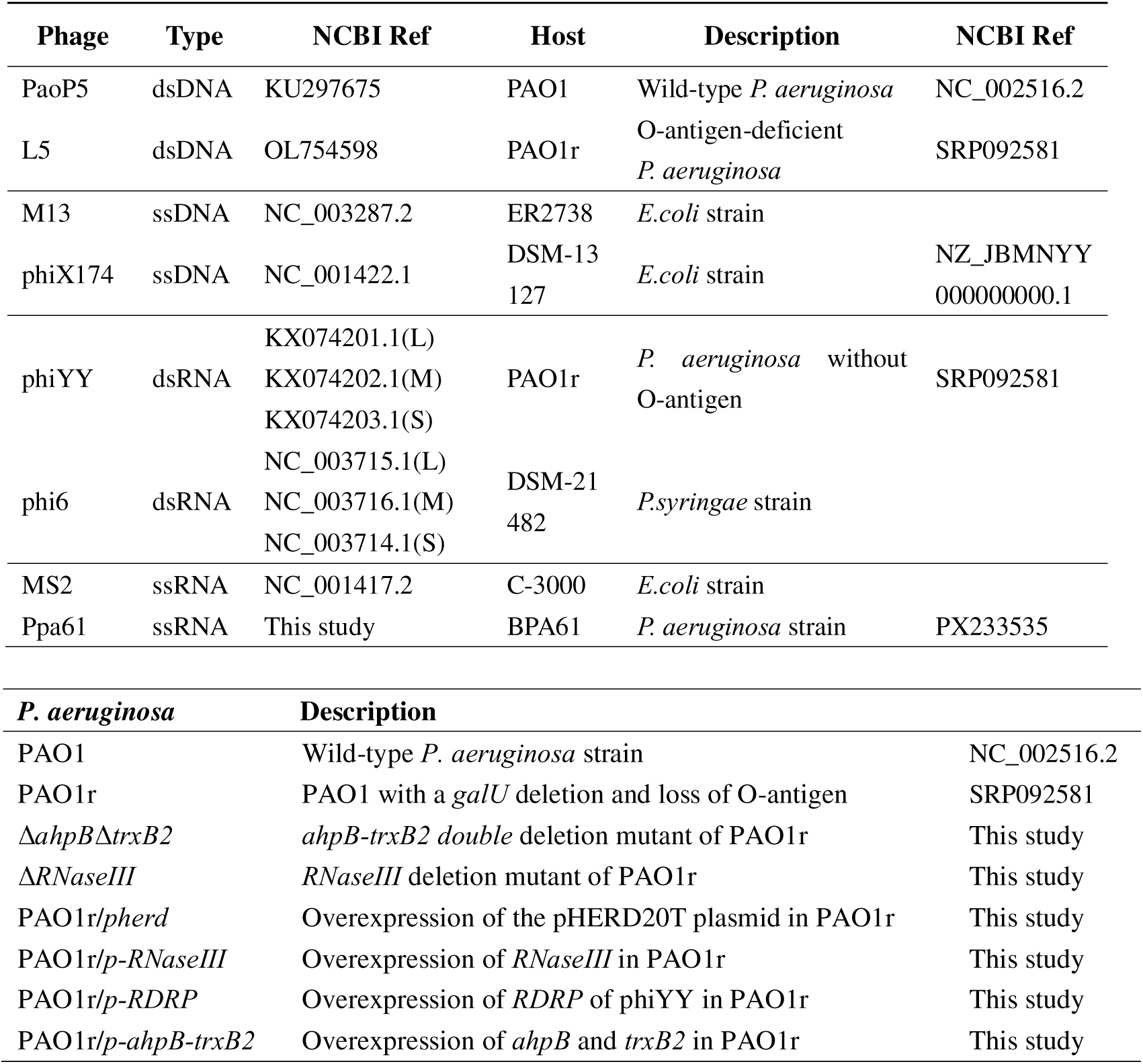
Bacterial strains and phages used in this study.

**TABLE S2:**
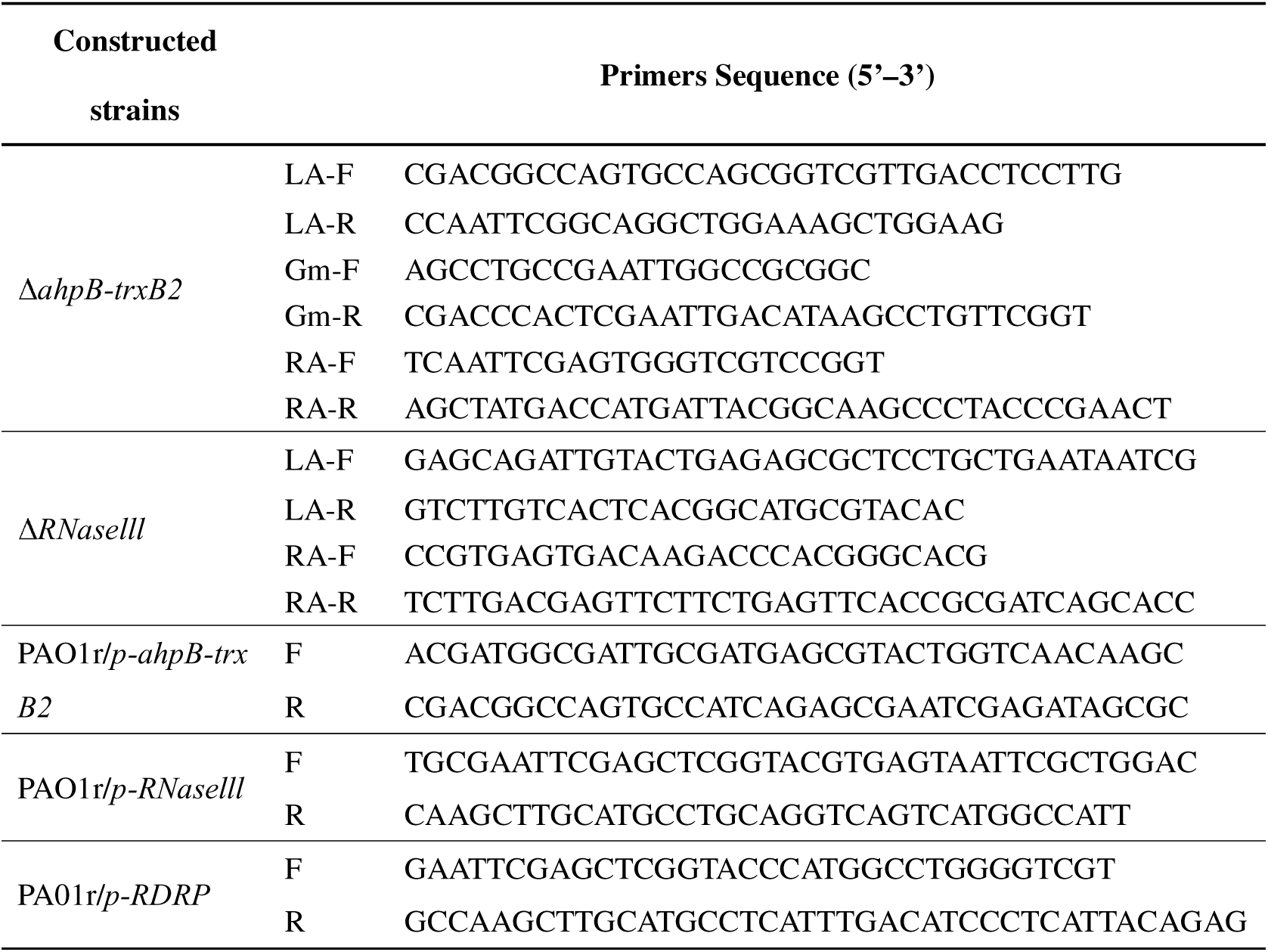
Primers used in this study.

**Fig. S1.**
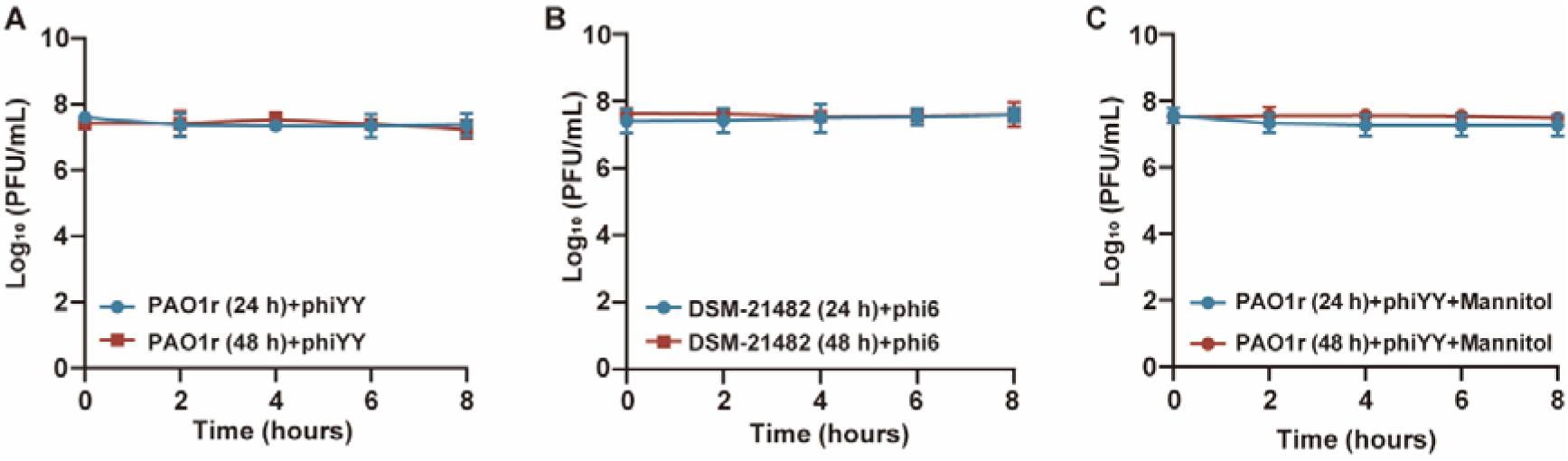
RNA phage survival dynamics. (A-B) Phage titers were monitored over 8 hours following infection of host bacteria cultured for 24 or 48 hours. phiYY showed no significant titer change in PAO1 cultures (A), and similarly, phi6 showed no significant change in DSM-21482 cultures (B), ruling out phage replication in these dormant-state cells. (C) To test whether mannitol reactivates host metabolism, phiYY was added to 24- or 48-hour PAO1 cultures pre-supplemented with 230 mM mannitol. No significant increase or decrease in phage titer was observed over 8 hours, indicating that mannitol does not disrupt bacterial dormancy or enable phage replication. Data are presented as mean ± s.d. (n = 3).

**Fig. S2:**
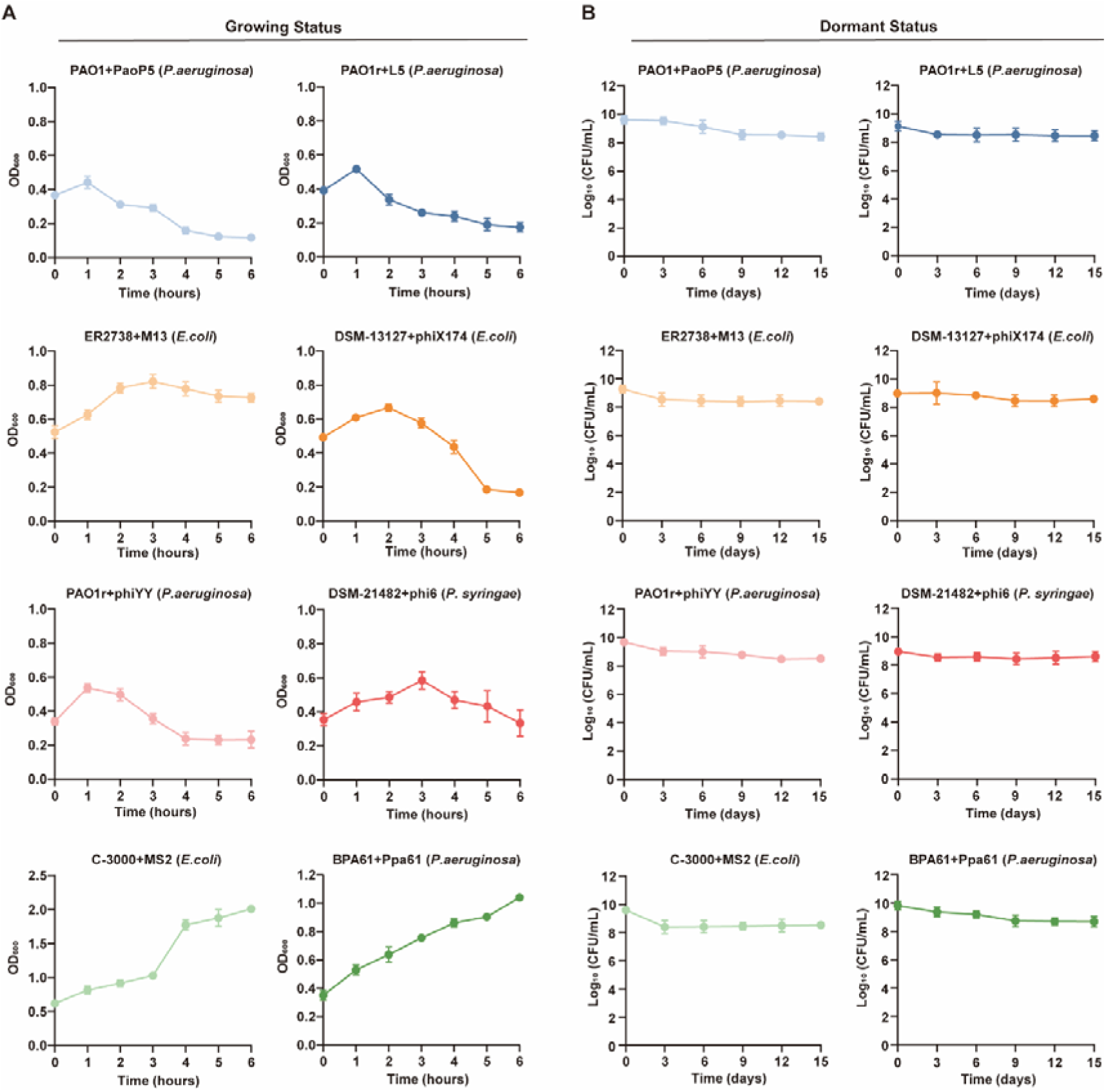
Bacterial survival dynamics in phage-host interactions. (A) Bacterial dynamics in log-phase cultures. Each of the eight phages was used to infect log-phase bacterial cultures at the indicated MOI. Bacterial growth was monitored by measuring optical density at 600 nm (OD_₆₀₀_) over time. (B) Bacterial dynamics in dormant cultures. Bacteria were cultured for 24 hours to induce dormancy prior to phage infection. Phages were then added at a multiplicity of infection (MOI) of 0.01. Bacterial viability was assessed at the indicated time points by serial 10-fold dilution and plating for colony-forming unit (CFU) enumeration. All experiments were performed in triplicate (n = 3), and data are presented as mean ± s.d..

